# The mechanism of Hsa_circ_0000629 in bronchial asthma through sponge adsorption of miR-212-5p/NLRP3

**DOI:** 10.64898/2026.03.21.713317

**Authors:** Su Xiandu, Lin Lieju, Yu Li, Guo Zhenli, Lin Mingli, Zeng Guidan, Chen Xu, Li Dongyang

## Abstract

To explore the mechanism of Hsa_circ_0000629 adsorbing miR-212-5p/ nucleotide-binding oligomerization domain-like receptor protein 3 (NLRP3) through sponge in bronchial asthma. Twenty BALB/C mice were randomly divided into a normal control group and an asthma group. Pathological changes in lung tissue were observed via HE staining. Human bronchial epithelial cells (16HBE) were transfected with Hsa_circ_0000629 overexpression group (Hsa_circ_0000629-over), Hsa_circ_0000629 siRNA (Hsa_circ_0000629-si), mimic NC, miR-212-5p mimic, inhibitor NC, miR-212-5p inhibitor, and LPS+Hsa_circ_0000629 si. LPS-induced asthmatic cell models (LPS group) and untransfected 16HBE cells (NC group) served as controls. qRT-PCR was used to measure Hsa_circ_0000629, miR-212-5p and NLRP3 expression. ELISA assessed interleukin 18 (IL-18), interleukin 1β (IL-1β), interleukin 6 (IL-6) and tumor necrosis factor -α (TNF-α) levels. Cell proliferation and the apoptosis were evaluated by EDU assay and flow cytometry, respectively. Western blot analyzed Cleaved-caspase 1, 3 and 9 proteins expression. Dual-luciferase assay verified the binding sites of Hsa_circ_0000629 to miR-212-5p and NLRP3 to miR-212-5p. HE staining revealed inflammatory cell infiltration, bronchial wall thickening, smooth muscle hyperplasia, and alveolar destruction in asthmatic mice. Compared with the controls, Hsa_circ_0000629 and NLRP3 expression were significantly increased, while miR-212-5p expression was decreased in asthmatic lung tissues. In 16HBE cells, Hsa_circ_0000629-over and LPS groups showed elevated Hsa_circ_0000629 and NLRP3 expression but reduced miR-212-5p levels. Silencing Hsa_circ_0000629 in LPS-treated cells (LPS+Hsa_circ_0000629-si) reversed these effects. Overexpression of miR-212-5p counteracted Hsa_circ_0000629-induced NLRP3 upregulation, while miR-212-5p inhibition enhanced NLRP3 expression. LPS exposure increased TNF-α, IL-18, IL-6, and IL-1β levels, reduced cell proliferation, and promoted apoptosis. These changes were attenuated by Hsa_circ_0000629 silencing or miR-212-5p overexpression. Western blot confirmed that Hsa_circ_0000629 overexpression upregulated Cleaved-Caspase 1, 3, and 9, whereas miR-212-5p mimic or Hsa_circ_0000629-si reversed this trend. Dual-luciferase assays demonstrated targeted interactions among Hsa_circ_0000629, miR-212-5p, and NLRP3. Interference with Hsa_circ_0000629 expression can alleviate LPS induced apoptosis in 16HBE cells and inhibit the expression of inflammatory factors by targeting the miR-212-5p/NLRP pathway, which may be a new target for the treatment of asthma.

## Introduction

Asthma is a prevalent chronic respiratory disease in children, characterized primarily by airway inflammation, airway remodeling, and heightened airway hyperresponsiveness. It has a high incidence rate and has become one of the major chronic diseases affecting children’s health worldwide^[1]^. Although current therapeutic agents have improved the prognosis for asthma patients, a subset of patients still responds poorly to treatment. Therefore, understanding the regulatory mechanisms underlying asthma pathogenesis is of considerable significance for the early prevention and treatment of this disease. Airway inflammation and airway remodeling are key factors driving the onset and progression of asthma.Nucleotide-binding oligomerization domain-like receptor protein 3 (NLRP3), as a key inflammasome, not only triggers airway inflammation by activating the inflammatory cytokine IL-1β but also cooperates with other signaling pathways to further exacerbate pyroptosis of airway epithelial cells, thereby contributing to the progression of asthma^[2]^. MicroRNA-212-5p (miR-212-5p), as a key regulator of gene expression, is involved in airway inflammatory responses and airway remodeling, and exerts protective effects against inflammation and oxidative stress, potentially playing a critical role in ameliorating the pathogenesis of asthma^[3]^. It has been reported that the miR-212-5p/NLRP3 signaling axis can induce chronic inflammation and oxidative stress, exacerbate inflammatory responses, and activate structural changes in the airways, thereby further aggravating airway inflammatory diseases^[4]^. In recent years, circular RNAs (hsa_circRNAs) in peripheral blood have garnered increasing attention and research interest, as they play a crucial role in respiratory diseases by modulating the expression of various inflammatory genes^[5]^. Studies have found that hsa_circ_0000629 may function as a miRNA sponge, modulating cellular immunity and inflammatory responses, thereby participating in the pathogenesis and progression of asthma, and may serve as a novel therapeutic target for asthma prevention and treatment^[6]^. However, it remains unclear whether hsa_circ_0000629 exerts its effects on asthma by directly targeting and regulating the miR-212-5p/NLRP3 axis.This study employs both animal and cellular experiments to investigate how hsa_circ_0000629 regulates NLRP3 expression by acting as a sponge for miR-212-5p, aiming to provide novel insights into the pathogenesis of asthma and pave the way for personalized clinical therapeutic strategies.

## Materials and Methods

Twenty specific pathogen-free (SPF) male BALB/C mice (8–9 weeks old, 22 ± 2 g) were obtained from Guangdong Zhiyuan Biomedicine Technology Co., Ltd. (License No.: SYXK (Yue) 2022-0301). All animal procedures were approved by the Animal Experiment Ethics Committee (Approval No.: MIS2023040).

### Experimental Animal

The mice were randomly divided into a normal control group and an asthma model group (asthma group), with 10 mice in each group. All mice were maintained under standard conditions with free access to food and water. For the asthma model group, an intraperitoneal injection of 0.2 mL of sensitization solution (containing 20 µg of grade V ovalbumin (9006-59-1, Psaitong) and 2 mg of aluminum hydroxide) was administered on days 0, 7, and 14. Starting from day 21, the mice were exposed to an aerosolized 2% grade II ovalbumin challenge solution using a compressed air-driven nebulizer. The challenges were performed three times per week, each lasting 30 minutes, for a total duration of 8 weeks. For the normal control group, sensitization and challenge procedures were replaced with an equal volume of normal saline at the corresponding time points.

### Cell Culture and Transfection

Human bronchial epithelial cells (16HBE cells, ORC0857, Shuangquan Biological Co., Ltd., Guangzhou) were cultured in a dedicated 16HBE medium (ORCCM0249, Yingxin Bio, Guangzhou) medium supplemented with 10% fetal bovine serum and 100 U/mL streptomycin-penicillin, and maintained in an incubator at 37°C with 5% CO_2_. The culture medium was refreshed every 2–3 days. Upon reaching 60–70% confluence, cells were transfected using Lipofectamine 2000 transfection reagent (SJ-LV-001-03, Yingxin Bio) according to the manufacturer’s protocol. The following groups were established: the Hsa_circ_0000629 overexpression group (Hsa_circ_0000629-over, using the Hsa_circ_0000629 overexpression vector, G70095-1, Shuangquan Biological Co., Ltd., Guangzhou), the Hsa_circ_0000629 interference group (Hsa_circ_0000629 siRNA, RY25006348, Shuangquan Biological Co., Ltd., Guangzhou; Hsa_circ_0000629-si), the negative control for the interference vector (mimic NC group), the miR-212-5p mimic group (miR-212-5p mimic), the negative control plasmid of the inhibitor (inhibitor NC group), the miR-212-5p inhibition group (miR-212-5p inhibitor), and the LPS + Hsa_circ_0000629-si group. The lipopolysaccharide (LPS)-induced asthmatic cell model was designated as the LPS group, while untransfected 16HBE cells were labeled as the blank control group (NC group).

### Hematoxylin and Eosin Staining of Mouse Lung Tissue

Mouse lung tissues were fixed in 4% paraformaldehyde for 48 hours, followed by dehydration, clearing, paraffin embedding, and sectioning (4 μm). For hematoxylin and eosin (HE) staining, the sections were first deparaffinized in xylene and rehydrated through a descending ethanol series to distilled water. Subsequently, the nuclei were stained with hematoxylin for 20 minutes, followed by differentiation and bluing. The cytoplasm was then counterstained with eosin for 3 minutes. Finally, the stained sections were dehydrated, cleared, and mounted with a neutral resinous mounting medium. Histopathological changes in the lung tissues were examined and imaged under a light microscope (SOPTOP).

### RT-qPCR Assay

Total RNA was extracted from mouse lung tissue using the Trizol reagent (P424, TIANGEN). Reaction system: 0.4 μ L of forward and reverse primers, 2 μ L of cDNA template, Hieff ™ Quantitative PCR SYBR ® Green premix (11201ES08, Yisheng Biotech) 10 μ L, double distilled water 10 μ L. The amplification conditions were: pre denaturation at 95 ℃ for 5 minutes, denaturation at 95 ℃ for 10 seconds, annealing at 60 ℃ for 30 seconds, and extension at 72 ℃ for 60 seconds, for a total of 40 cycles. Calculate the content of Hsa_circ_0000629 (GAPDH internal reference), miR-212-5p (U6 internal reference), and NLRP3 (U6 internal reference) using the 2^-△△Ct^ method.

### Enzyme-Linked Immunosorbent Assay

The concentrations of TNF-α, IL-18, IL-6, and IL-1 β in cell culture supernatants were quantified using commercial enzyme-linked immunosorbent assay (ELISA) kits (LunChangShuo Biotech, China) according to the manufacturer’s instructions. Briefly, collected cells were washed three times with PBS, and 1–2 × 10^6^ cells were resuspended in 150–200 μL of PBS. Cells were lysed by repeated freeze-thaw cycles. The lysate was then centrifuged at 2000 rpm for 10 minutes, and the supernatant was collected for analysis. After adding standard or sample solutions (50 μl) to the assigned wells, 50 μl detection reagent were added to all wells, and then add 100 μl of HRP-conjugate reagent to each well, cover with an adhesive strip and incubate for 45 min at 37 °C. After washing 5 times with 400 μl wash solution, 50 μl chromogen solution A and 50 μl chromogen solution B were added to each well to incubate for 15 min at 37 °C. The reaction was stopped by adding 50 μl stop solution. The absorbance was immediately measured at 450 nm using a microplate reader.

### EDU detection of cell proliferation experiment

After overexpression or interference transfection with Hsa_circ_0000629, 16HBE cells were digested, counted, and seeded in 96 well plates with 5000 cells per well (3 replicates per group). After cell adhesion, add culture medium containing 50 μ M EdU and incubate for 2 hours to label proliferating cells. Subsequently, discard the culture medium, wash twice with PBS, fix with 4% paraformaldehyde for 30 minutes, and quench with glycine for 5 minutes; After washing with PBS and cell infiltration treatment, add EdU reaction solution (Guangzhou Shuangquan Biological) and react in the dark for 30 minutes. Finally, stain the nuclei with Hoechst for 30 minutes. Evaluate cell proliferation by taking pictures under specific excitation conditions (Hoechst: 350 nm/50 ms; EdU: 567 nm/500 ms) using a fluorescence microscope.

### Flow cytometry Assay

After 48 hours of cell culture in each group, the cells were digested with trypsin, collected by centrifugation, washed with pre cooled PBS, and resuspended in 1 × Binding Buffer (1 × 10 ^6^ cells/mL). Take 100 µ L of cell suspension, add 5 µ L of FITC Annexin V and 5 µ L of PI, incubate at room temperature in the dark for 15 minutes, then add 400 µ L of 1×Binding Buffer, and detect cell apoptosis by flow cytometry (D2060R.Agilent, Novocyte).

### Western blot

The cell proteins were extracted from each group, 35 μg of proteins were separated on 10% SDS-PAGE gels and transferred onto PVDF membranes. Protein membranes were pretreated with skim milk for 2 h at room temperature and then incubated with antibodies against target proteins at 4 °C overnight. The primary antibodies included Cleaved Caspase 1 (AF4005, Affinity), Cleaved Caspase 3 (AF7022, Affinity), and Cleaved Caspase 9 (AF5240, Affinity). Loading controls were using antibodies against β-actin (T0023, Affinity). The protein membranes were further incubated with HRP-conjugated secondary antibodies. Protein bands were detected using an enhanced chemiluminescence substrate and visualized with an automated chemiluminescence image system (LAS-3000, FUJIFILM).

### Dual luciferase assay

Cells were transfected using cationic liposome method, and dual luciferase activity was detected according to the instructions of the dual luciferase reporter gene detection kit (RG027, Beyotime).

### Data Analysis

Statistical analysis was performed using SPSS 27.0 software. Normally distributed measurement data were expressed as (mean ± SD). Comparisons between two groups were conducted using the t-test, while comparisons among multiple groups were analyzed by one-way ANOVA, followed by the LSD-t test for pairwise comparisons. *P* < 0.05 was considered statistically significant.

## Result

### HE staining for detecting pathological conditions of lung tissue

To assess the pathological features of the asthma model, lung tissue sections were examined. Compared with the normal control group, lung tissues from asthmatic mice exhibited significant inflammatory cell infiltration, characterized by extensive accumulation of eosinophils around the bronchi and in peribronchial regions. Additionally, there was notable thickening of the bronchial walls and smooth muscle hyperplasia. Alveolar destruction was also evident, including thickened alveolar septa and collapsed or fused alveolar spaces (Fig.1). These collective pathological alterations confirm the successful establishment of the murine asthma model, and are consistent with the results of previous studies^[4]^.

**Figure 1.**
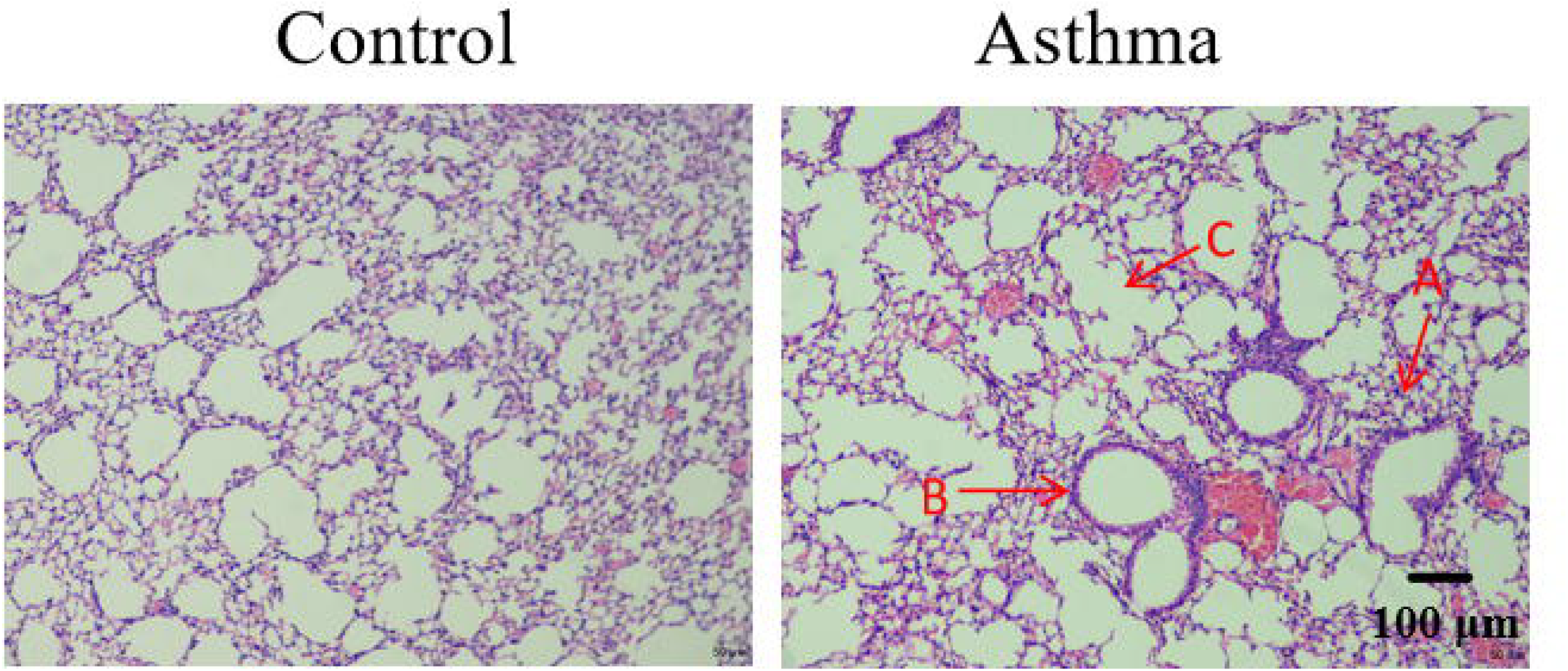
HE staining of mouse lung tissue. A, Inflammation cell infiltration, with a large number of eosinophils accumulating in the bronchi and surrounding areas. B, Bronchial wall thickening and smooth muscle hyperplasia. C, Thickening of alveolar septa, collapse or fusion of alveolar cavities.

### Expression and Regulation of the Hsa_circ_0000629–miR-212-5p–NLRP3 Axis in an Asthma Model

In asthma, NLRP3 serves as a critical component of the inflammasome and is involved in airway inflammation and pyroptosis. miR-212-5p may influence this process by targeting and regulating NLRP3 or related pathway genes. The circular RNA Hsa_circ_0000629 potentially acts as a sponge for miR-212-5p^[6]^, thereby indirectly modulating the expression of downstream target genes and participating in the inflammatory regulatory mechanisms of asthma. Consequently, its expression was examined in lung tissues from an asthma model group. As Figure 2 A shows, the expression levels of Hsa_circ_0000629 and NLRP3 were significantly increased (p < 0.001), while miR-212-5p expression was markedly decreased (p < 0.001) in lung tissues of the asthma group. Similarly, we also measured its expression level in cell experiments.

**Figure 2.**
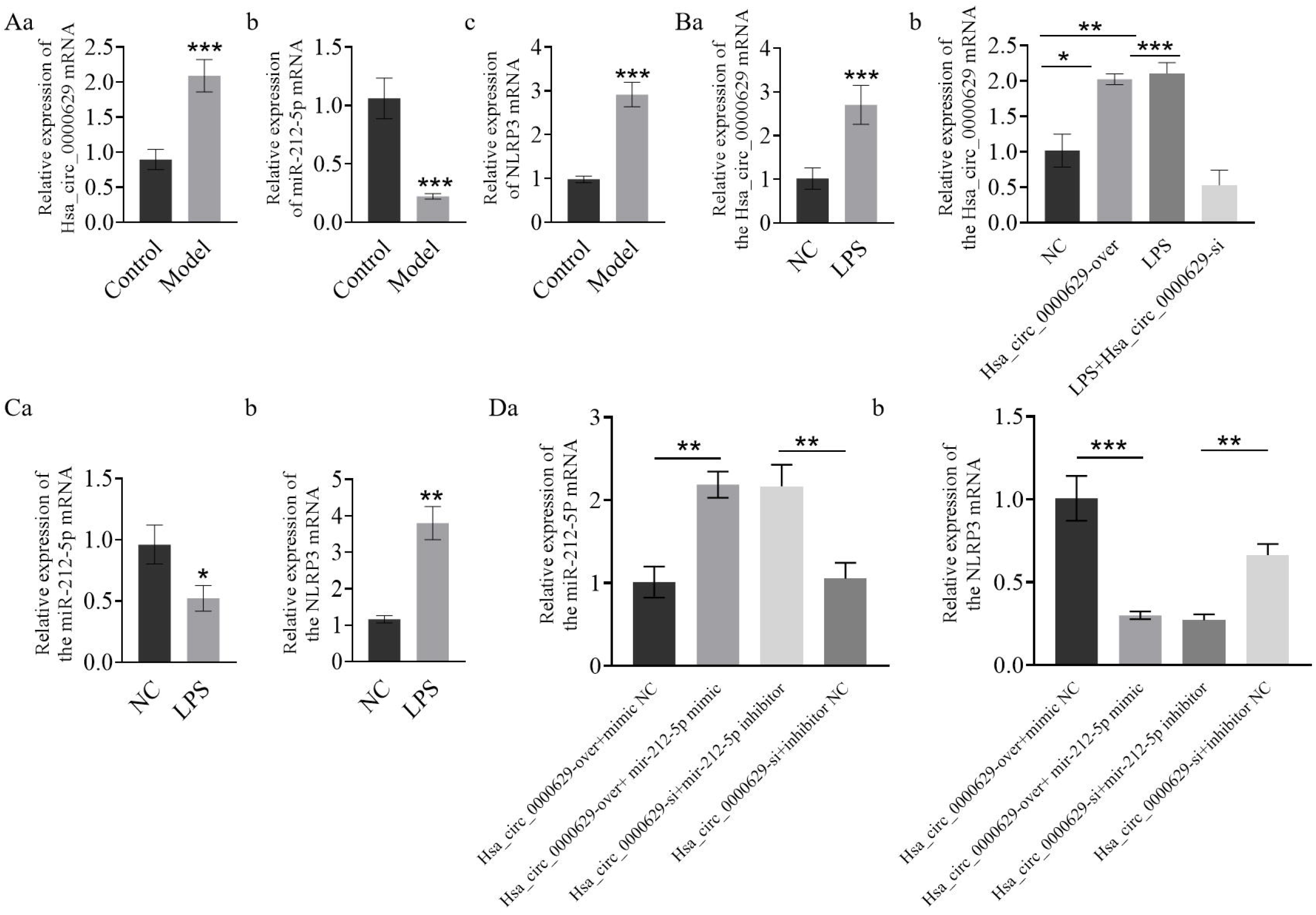
Expression levels of Hsa_circ_0000629, miR-212-5p, and NLRP3 in asthma model and cells. A, The expression levels of Hsa_circ_0000629, miR-212-5p and NLRP3 in the asthma model group and the control group. B,The expression levels of Hsa_circ_0000629 in each group of cells. C, The expression levels of miR-212-5p and NLRP3 in the NC group and the LPS group. D,The expression levels of miR-212-5p and NLRP3 in each group of cells. **, P < 0.01, ***, P < 0.001, vs. control. Abbreviations: LPS, lipopolysaccharide.

The expression level of Hsa_circ_0000629 was significantly higher in the Hsa_circ_0000629-over group and the LPS group compared to the NC group (p < 0.05 and p < 0.01, respectively). The expression level of Hsa_circ_0000629 in the LPS + Hsa_circ_0000629-si group was significantly lower than that in the LPS group (p < 0.001, Fig. 2B). Compared with the NC group, the expression level of miR-212-5p in the LPS group was significantly decreased (p < 0.05), while the expression of NLRP3 was significantly increased (p < 0.01, Fig. 2C). Compared with the Hsa_circ_0000629 over+mimic NC group, the expression level of miR-212-5p in the Hsa_circ_0000629 over+miR-212-5p mimic group was significantly increased (p < 0.001), while the expression of NLRP3 was significantly decreased (p < 0.001, Fig. 2D). Compared with the Hsa_circ_0000629-si+inhibitor NC group, the expression level of miR-212-5p in the Hsa_circ_0000629-si+miR-212-5p inhibitor group was significantly decreased (p < 0.001), while the expression of NLRP3 was significantly increased (p < 0.001, Fig. 2D).This indicates that sa_circ_0000629 regulates NLRP3 expression through miR-212-5p.

### To validate the expression levels of inflammatory factors at the cellular level

To validate the expression levels of inflammatory factors following asthma, we utilized an LPS-induced asthma cell model for analysis. The results demonstrated that, compared with the control group, the levels of TNF - α, IL-18, IL-6, and IL-1 β were significantly increased in the LPS group (p < 0.01, Fig. 3).These findings confirm at the cellular level that LPS stimulation effectively mimics the key inflammatory state in asthma, particularly by activating the inflammatory pathway centered on the NLRP3 inflammasome and accompanied by pyroptosis, thereby providing a reliable model basis for studying the mechanisms of airway inflammation in asthma.

**Figure 3.**
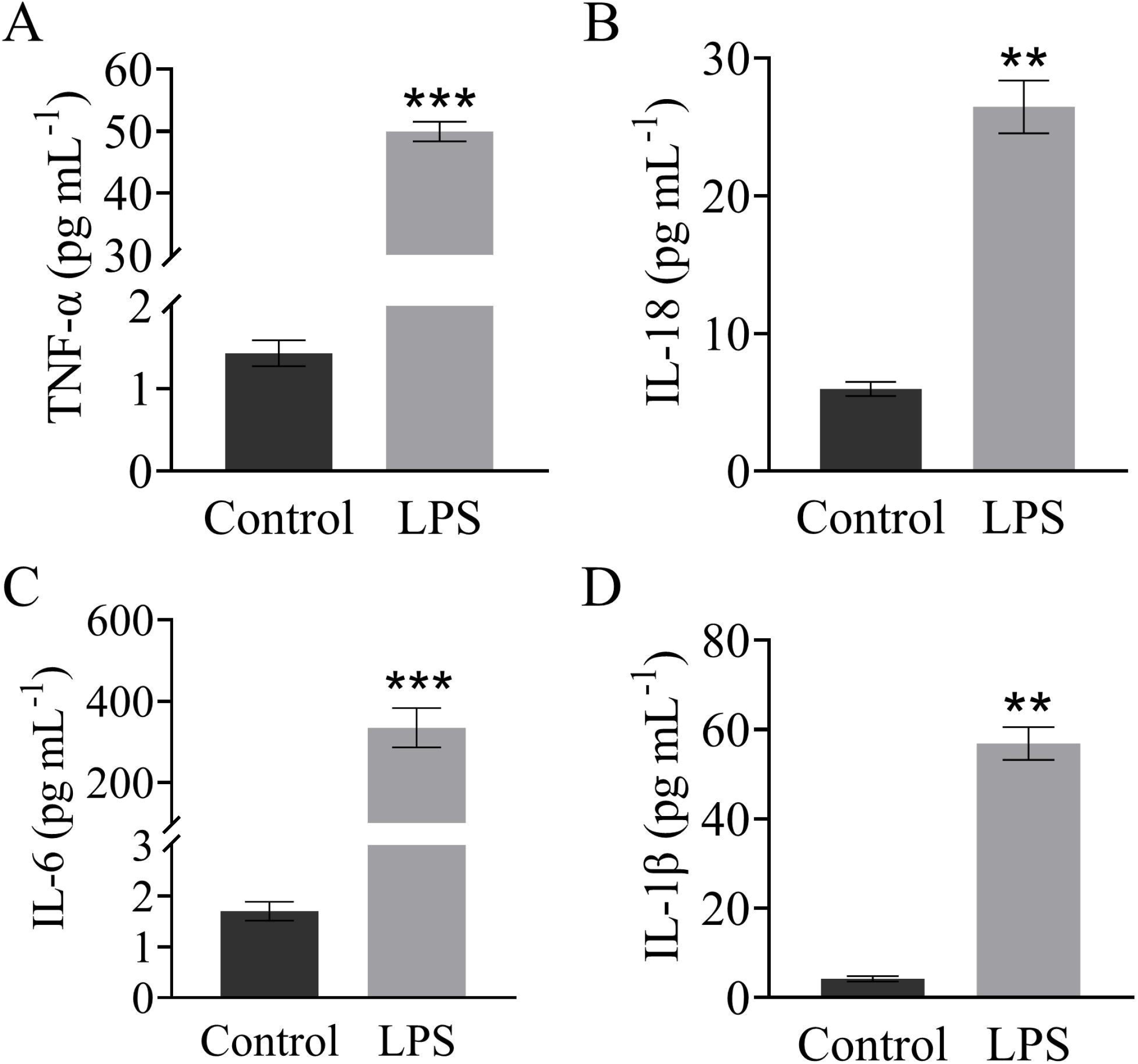
Comparison of TNF - α, IL-18, IL-6, and IL-1 β levels between LPS group and control. group. A, Comparison of TNF-α levels between the LPS group and the control group. B, Comparison of IL-18 levels between the LPS group and the control group. C, Comparison of IL-6 levels between the LPS group and the control group. D, Comparison of IL-1β levels between the LPS group and the control group. **, P < 0.01, ***, P < 0.001, vs. control. Abbreviations: LPS, lipopolysaccharide.

### The effect of overexpression or interference of Hsa_circ_0000629 on the proliferation ability of 16HBE cells

To understand the regulatory effect of Hsa_circ_0000629 on the proliferation ability of 16HBE cells, Compared with the NC group, the proliferation rate of 16HBE cells in the LPS group and Hsa_circ_0000629 over group was significantly reduced (p < 0.001). The proliferation rate of 16HBE cells in the LPS+Hsa_circ_0000629-si group was significantly higher than that in the LPS group(p < 0.001). Compared with the Hsa_circ_0000629 over+mimic NC group, the proliferation rate of 16HBE cells in the Hsa_circ_0000629 over+miR-212-5p mimic group was significantly increased (p < 0.001). Compared with the Hsa_circ_0000629-si+inhibitor NC group, the proliferation rate of 16HBE cells in the Hsa_circ_0000629-si+miR-212-5p inhibitor group was significantly reduced (p < 0.001, Figure 4). This indicates that restoring the expression of miR-212-5p can reverse the changes in proliferation ability caused by alterations in the expression level of Hsa_circ_0000629.

**Figure 4.**
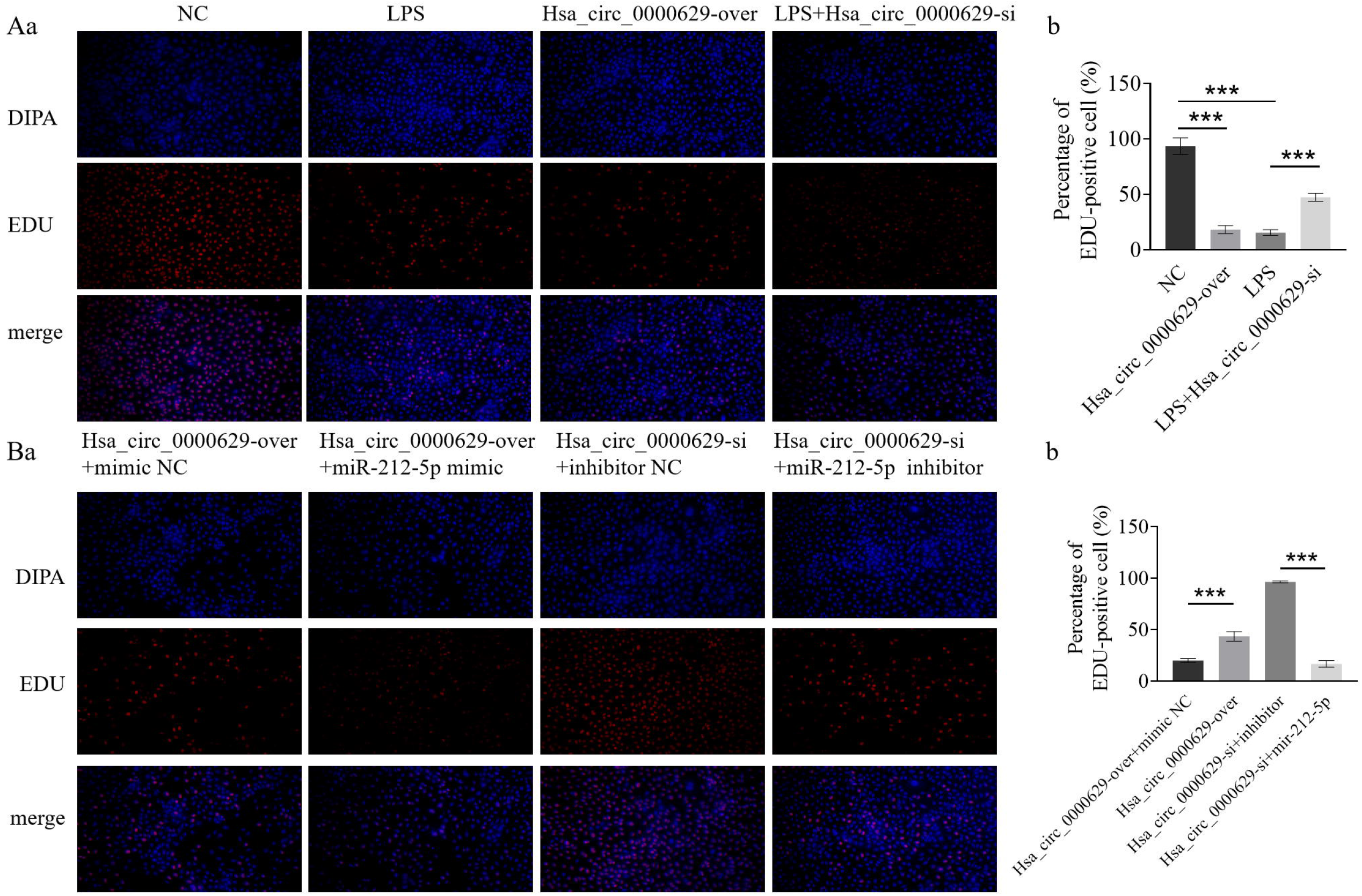
Poliferation of 16HBE cells in each group. A, Cell proliferation in LPS group and blank control group. B, Cell proliferation in Hsa_circ_0000629-over group and LPS+Hsa_circ_0000629-si group. C, Cell proliferation after overexpression and inhibition of miR-212-5p in Hsa_circ_0000629-over group and Hsa_circ_0000629-si group.**, P < 0.01, ***, P < 0.001, vs. control. Abbreviations: LPS, lipopolysaccharide.

### The effect of overexpression or interference of Hsa_circ_0000629 on the apoptosis ability of 16HBE cells

To clarify the regulatory effect of Hsa_circ_0000629 on apoptosis in 16HBE cells, Compared with the NC group, the apoptosis rate of 16HBE cells in the Hsa_circ_0000629 over group was significantly increased (p < 0.001), while the apoptosis rate of 16HBE cells in the Hsa_circ_0000629 si group was significantly decreased (p < 0.001). The apoptosis rate of 16HBE cells in the LPS+Hsa_circ_0000629-si group was significantly lower than that in the LPS group (p < 0.001). Compared with the Hsa_circ_0000629 over+mimic NC group, the apoptosis rate of 16HBE cells in the Hsa_circ_0000629 over+miR-212-5p mimic group was significantly reduced (p < 0.001). Compared with the Hsa_circ_0000629-si+inhibitor NC group, the apoptosis rate of 16HBE cells in the Hsa_circ_0000629-si+miR-212-5p inhibitor group was significantly increased (p < 0.001, Figure 5). This indicates that the changes in cell apoptosis rate induced by altered Hsa_circ_0000629 expression levels can be reversed.

**Figure 5.**
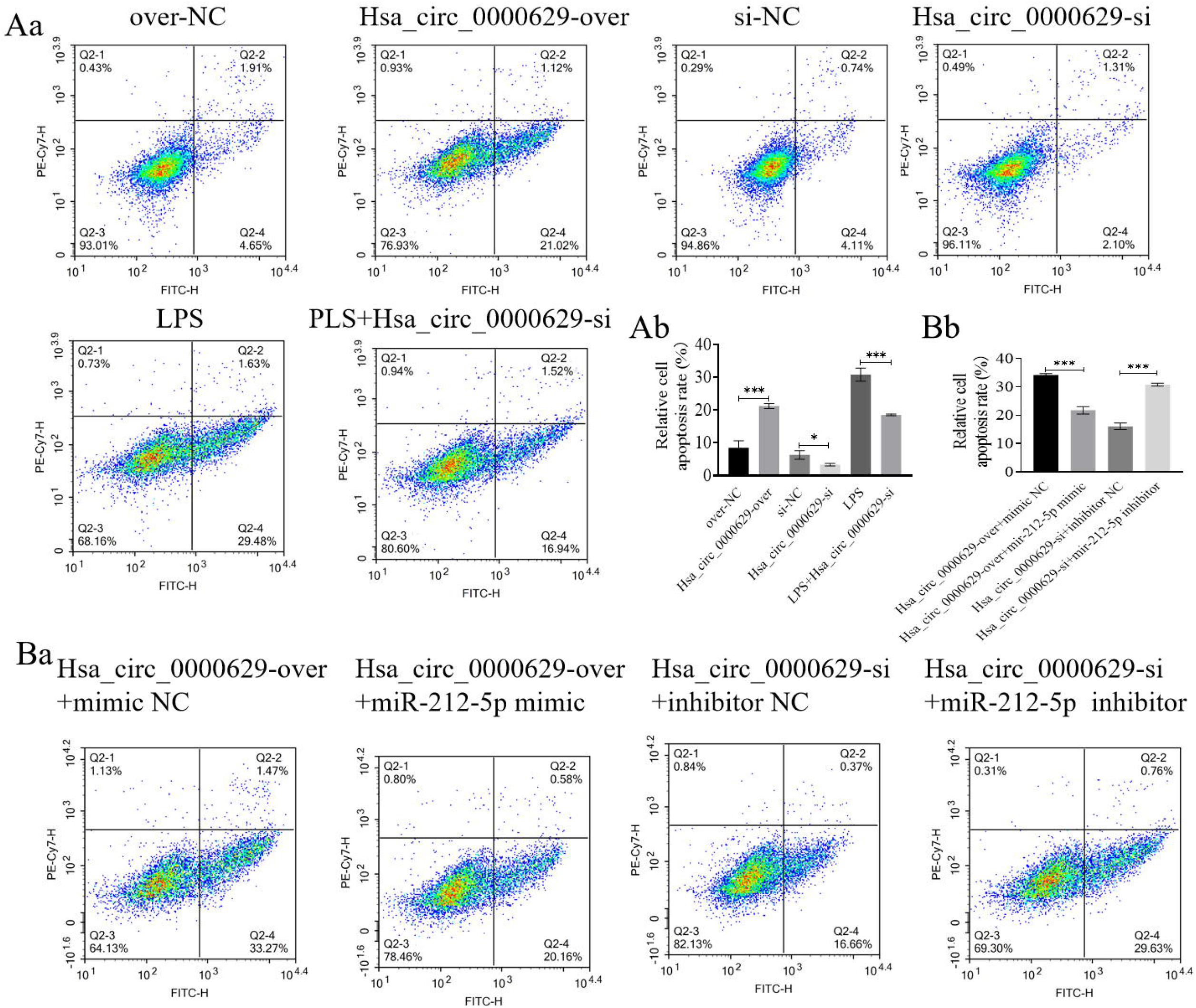
Apoptosis of 16HBE cells in each group. A, Apoptosis rate of 16HBE cells in blank control group, LPS group, Hsa_circ_0000629 over group, and LPS+Hsa_circ_0000629 si group. B, Apoptosis rate of 16HBE cells after overexpression and inhibition of miR-212-5p in Hsa_circ_0000629 over group and Hsa_circ_0000629 si group.**, P < 0.01, ***, P < 0.001, vs. control. Abbreviations: LPS, lipopolysaccharide.

### Western blot detection of protein expression levels of Cleaved Caspase 1, 3, and 9

By detecting the protein expression levels of Cleaved-Caspase 1, 3, and 9 in the cells, the activation degree of cell apoptosis under different experimental conditions was evaluated, thereby exploring the mechanism of cell death.Compared with the NC group, the expression levels of Cleaved Caspase 1, 3, and 9 proteins were significantly increased in the Hsa_circ_0000629 over group (p < 0.001, p < 0.001, p < 0.01). The expression levels of Cleaved Caspase 1, 3, and 9 proteins in the LPS+Hsa_circ_0000629-si group were significantly lower than those in the LPS group(p < 0.001). Compared with the Hsa_circ_0000629 over+mimic NC group, the expression levels of Cleaved Caspase 1, 3, and 9 proteins in the Hsa_circ_0000629 over+miR-212-5p mimic group were significantly decreased (p < 0.001). Compared with the Hsa_circ_0000629-si+inhibitor NC group, the expression levels of Cleaved Caspase 1, 3, and 9 proteins were significantly increased in the Hsa_circ_0000629-si+miR-212-5p inhibitor group (p < 0.001, Figure 6).

**Figure 6.**
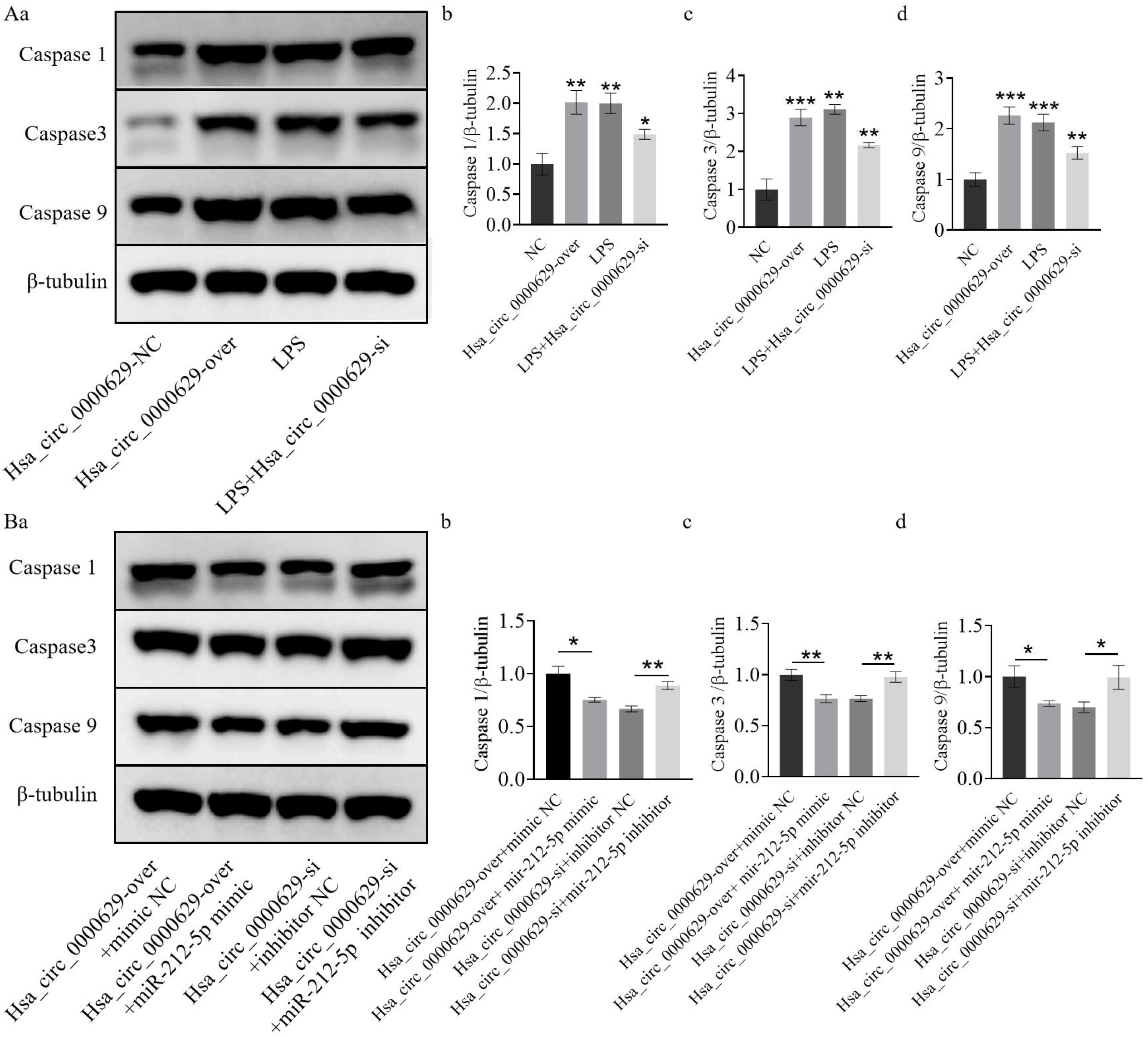
Faved Caspase1, 3, and 9 proteins are expressed in the Hsa_circ_0000629-over group and Hsa_circ_0000629-si group.

### 2.8 Dual luciferase assay

By detecting the changes in luciferase activity, it was confirmed whether there was a targeted regulatory relationship among Hsa_circ_0000629, miR-212-5p and NLRP3.Dual-luciferase reporter assays showed that, compared with the Hsa_circ_0000629-WT vector co-transfected with miR-212-5p mimic, the Hsa_circ_0000629-MUT vector co-transfected with miR-212-5p mimic exhibited significantly increased reporter fluorescence (p < 0.001, Fig. 7A). Conversely, compared with the Hsa_circ_0000629-WT vector co-transfected with miR-212-5p inhibitor, the Hsa_circ_0000629-MUT vector co-transfected with miR-212-5p inhibitor showed significantly decreased reporter fluorescence (p < 0.001, Fig. 7A). These results indicate that miR-212-5p directly binds to its complementary site on Hsa_circ_0000629 to regulate reporter gene expression in this experimental model.

**Figure 7.**
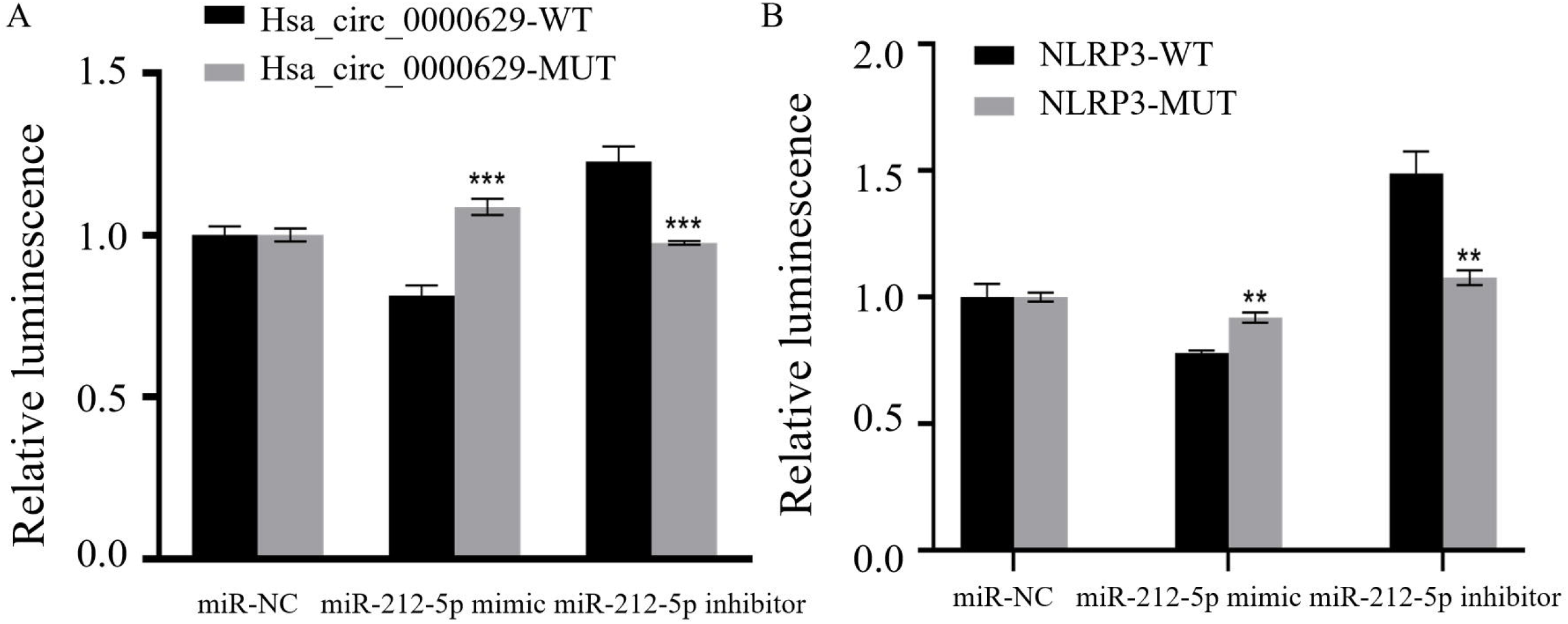
Expression analysis of dual luciferase reporter genes. A, Hsa_circ_0000629 and miR-212-5pN. B,NLRP3 and miR-212-5p.

Compared with the miR-212-5p mimic group of NLRP3-WT vector, the reported fluorescence of miR-212-5p mimic in NLRP3-MUT vector was significantly increased (p < 0.01, Fig. 7B); Compared with the NLRP3-WT vector miR-212-5p inhibitor, the NLRP3-MUT vector miR-212-5p inhibitor reported a significant decrease in fluorescence (p < 0.01, Fig. 7B). The expression analysis of dual luciferase reporter genes showed that miR-212-5p mimic had a significant downregulation effect on the reporter fluorescence of NLRP3-WT vector. After the corresponding binding site mutation, there was no significant change in the reporter fluorescence of the mutant vector. In this experimental model, miR-212-5p can regulate the expression of reporter genes through the corresponding binding sites on NLRP3.

## Discussion

The pathogenesis and etiology of asthma remain unclear. Most studies suggest that its development is primarily associated with airway inflammation, immune function, environmental factors, and genetic factors^[7-8]^. Asthma is characterized by episodes of bronchoconstriction, airway hyperresponsiveness, and remodeling. A significant consequence of persistent inflammation is damage to the pulmonary tissue, which can disrupt local neural architecture and signaling.The airway inflammation characterized primarily by eosinophil infiltration is an important reference indicator for judging the success of the asthma mouse model^[9]^.We observed characteristic pathological features in asthmatic mice, including bronchial wall thickening, smooth muscle hyperplasia, alveolar destruction, and inflammatory cell infiltration. Specifically, the accumulation of eosinophils (a key inflammatory cell type in asthma) was evident^[10]^ (Fig.1). Repeated epithelial injury triggers airway hyperresponsiveness via cascade reactions, a key mechanism in asthma onset and progression^[11]^. Within this complex inflammatory network, a key molecular hub, the NLRP3 inflammasome, has garnered widespread attention. NLRP3 is an inflammasome discovered in recent years, and its transcription gene is located on human chromosome 10 1q44, encoding a protein that plays an important role in airway inflammatory diseases^[12]^. Previous studies have shown that NLRP3 is not only closely related to the activation of airway inflammatory cells and related signaling pathways in asthma, but also mediates airway remodeling, exacerbates asthma symptoms, and participates in the progression of asthma disease^[13-14]^. In our study, we found a significant upregulation of NLRP3 expression at both asthmatic mice lung tissue and cells (Fig. 2C). In summary, the upregulation of NLRP3 is not only a hallmark of the inflammatory state in asthma, but also likely serves as a core molecular link driving airway hyperresponsiveness, remodeling, and disease progression.

MiR-212-5p is a multifunctional regulatory factor involved in cell growth, apoptosis, differentiation, and metabolism. Its overexpression can inhibit inflammatory responses and play a key role in lung tissue inflammation^[15]^. Consistent with this, Xu et al^[16]^ reported that miR-212-5p is downregulated in chronic lung diseases and can inhibit cigarette smoke–induced airway epithelial cell apoptosis, demonstrating a protective role in ameliorating apoptosis of human pulmonary microvascular endothelial cells and lung inflammation. The miR-212-5p/NLRP3 axis, as an important signaling pathway, participates in the pathogenesis and progression of airway diseases by inducing apoptosis and inflammatory responses in airway epithelial cells. Hsa_circ_0000629, a non coding RNA related to tumors, is involved in cell proliferation, metabolic regulation, and tissue differentiation.Its dysregulation of expression is not only involved in the pathogenesis of tumors, but also related to the body’s inflammatory and immune responses^[17]^. Bao et al ^[18]^found that Hsa_circ_0000629 is upregulated in the lung tissues of asthmatic mice and is involved in regulating inflammatory responses in airway epithelial cells. Our findings align with and extend these observations. In asthmatic lung tissue, we observed increased expression of Hsa_circ_0000629 and NLRP3, alongside decreased miR-212-5p expression. In vitro, compared with the NC group, the expression levels of Hsa_circ_0000629 and NLRP3 were significantly increased in the Hsa_circ_0000629 over group and LPS group, while the expression level of miR-212-5p was significantly decreased. These findings suggest that the upregulation of Hsa_circ_0000629 and NLRP3, along with the downregulation of miR-212-5p, may be involved in the pathogenesis of asthma. Further analysis of the transfected cells in each group revealed that, after LPS stimulation, the expression level of Hsa_circ_0000629 in the Hsa_circ_0000629 knockdown group was significantly lower than that in the LPS-only group. When miR-212-5p expression remained unchanged, altering Hsa_circ_0000629 expression alone did not affect NLRP3 expression. In the Hsa_circ_0000629-overexpression + miR-212-5p mimic group, miR-212-5p expression was increased and NLRP3 expression was decreased. Conversely, in the Hsa_circ_0000629-siRNA + miR-212-5p inhibitor group, miR-212-5p expression was reduced, accompanied by an increase in NLRP3 expression. These findings indicate that Hsa_circ_0000629, NLRP3, and miR-212-5p are involved in the pathogenesis and progression of asthma, and that Hsa_circ_0000629 may regulate NLRP3 expression through miR-212-5p. Furthermore, we measured key inflammatory cytokines (TNF-α, IL-18, IL-6, IL-1β) known to promote airway inflammation and hyperresponsiveness. Their levels were significantly elevated in the LPS group, suggesting their contribution to asthmatic airway inflammation. This aligns with reports that circRNAs can sponge asthma-related miRNAs to regulate inflammatory genes ^[19]^, and that miR-212-5p downregulation promotes disease progression^[20]^. Additionally, NLRP3 is known to exacerbate asthma by driving inflammatory cytokine expression and macrophage polarization ^[21]^(Fig.2, Fig.3).

In addition, we investigated the functional impact of Hsa_circ_0000629 on cellular processes. Overexpression of Hsa_circ_0000629 significantly increased the rate of cell apoptosis and markedly decreased cell proliferation, whereas knockdown of Hsa_circ_0000629 significantly reduced apoptosis and notably enhanced cell proliferation.Restoring miR-212-5p expression reversed the changes in cell proliferation caused by alterations in Hsa_circ_0000629 levels. These findings indicate that Hsa_circ_0000629 suppresses cell proliferation and promotes apoptosis, and that miR-212-5p acts as a key downstream effector molecule mediating the biological functions of Hsa_circ_0000629. Hsa_circ_0000629 likely exerts its effects by sequestering or sponging miR-212-5p, thereby modulating its activity. To elucidate the underlying apoptotic mechanism, we examined key executioner proteins. Caspase-1 amplifies inflammatory responses by activating inflammasome-mediated IL-1 β /IL-18 expression and inducing pyroptosis, thereby promoting allergic inflammation ^[22]^. Caspase-3 and Caspase-9 proteins primarily exacerbate airway injury by inducing epithelial cell apoptosis, disrupting airway barrier function, promoting inflammatory cell infiltration, and enhancing airway hyperresponsiveness^[23]^. Together, cleaved forms of these caspases form a core network governing cell death and inflammation in asthma. Consistent with the observed cellular phenotypes, the expression levels of cleaved Caspase-1, Caspase-3, and Caspase-9 were increased in the Hsa_circ_0000629-overexpression group. Conversely, their expression was reduced in the LPS + Hsa_circ_0000629-siRNA group compared to the LPS-only group (Fig.6A). Furthermore, in the Hsa_circ_0000629-overexpression + miR-212-5p mimic group, the levels of these three proteins were lower than those in the Hsa_circ_0000629-overexpression + mimic negative control(NC) group. Conversely, in the Hsa_circ_0000629-siRNA + miR-212-5p inhibitor group, the expression of the three proteins was higher than that in the Hsa_circ_0000629-siRNA + inhibitor NC group. These results indicate that Hsa_circ_0000629 promotes cell apoptosis by negatively regulating miR-212-5p expression, thereby influencing the activation of cleaved Caspase-1, Caspase-3, and Caspase-9. Finally, to directly validate the molecular interaction, Dual-luciferase reporter assays demonstrated that Hsa_circ_0000629 functions as a competing endogenous RNA by acting as a molecular sponge for miR-212-5p, thereby reducing the free intracellular concentration of miR-212-5p. This sequestration alleviates the repressive effect of miR-212-5p on its downstream target gene NLRP3, ultimately leading to upregulated NLRP3 expression. These findings confirm that both Hsa_circ_0000629 and NLRP3 are direct targets of miR-212-5p, which negatively regulates their expression by binding to specific complementary sequences within both molecules. This provides crucial molecular evidence for understanding the pathological mechanisms underlying asthma.

In summary, the expression levels of Hsa_circ_0000629 and NLRP3 are upregulated, while miR-212-5p is downregulated, in bronchial asthma tissues and cells. Silencing Hsa_circ_0000629 alleviates LPS-induced apoptosis in 16HBE cells and suppresses the expression of inflammatory cytokines by targeting the miR-212-5p/NLRP3 pathway, suggesting that Hsa_circ_0000629 may represent a novel therapeutic target for asthma. However, given the complexity of the mechanisms underlying asthma pathogenesis, it remains unclear whether other hsa_circular RNAs and signaling pathways also participate in regulating proliferation and apoptosis of 16HBE cells in asthma, warranting further in-depth investigation through additional basic experimental studies.

## Ethical approval and consent to participate

Protocols for animal experiments were approved by the Animal Experimental Ethics Committee of the Guangzhou Miles Biotechnology Co., Ltd. (Approval No.: MIS2023040), in compliance with the National Institutes of Health guidelines for the care and use of laboratory animals.

## Declaration of conflicting interests

The authors declared no potential conflicts of interest with respect to the research, authorship, and/or publication of this article.

## Notes

### Competing Interest Statement

The authors have declared no competing interest.

### Funding

- Hainan Provincial Health and Wellness Science and Technology Innovation Joint Project: WSJK2024MS202

